# MCRiceRepGP: a framework for identification of sexual reproduction associated coding and lincRNA genes in rice

**DOI:** 10.1101/271353

**Authors:** Agnieszka A. Golicz, Prem L. Bhalla, Mohan B. Singh

**Author notes:** Corresponding author: Agnieszka A. Golicz. Corresponding author: Mohan B. Singh.

## Abstract

Sexual reproduction in plants underpins global food production and evolution. It is a complex process, requiring intricate signalling pathways integrating a multitude of internal and external cues. However, key players and especially non-coding genes controlling plant sexual reproduction remain elusive. We report the development of MCRiceRepGP a novel machine learning framework, which integrates genomic, transcriptomic, homology and available phenotypic evidence and employs multi-criteria decision analysis and machine learning to predict coding and non-coding genes involved in rice sexual reproduction.

The rice genome was re-annotated using deep sequencing transcriptomic data from reproduction-associated tissues/cell types identifying novel putative protein coding genes, transcript isoforms and long intergenic non-coding RNAs (lincRNAs). MCRiceRepGP was used for genome-wide discovery of sexual reproduction associated genes in rice; 2,275 protein-coding and 748 lincRNA genes were predicted to be involved in sexual reproduction. The annotation performed and the genes identified, especially the ones for which mutant lines with phenotypes are available provide a valuable resource. The analysis of genes identified gives insights into the genetic architecture of plant sexual reproduction. MCRiceRepGP can be used in combination with other genome-wide studies, like GWAS, giving more confidence that the genes identified are associated with the biological process of interest. As more data, especially about mutant plant phenotypes will become available, the power of MCRiceRepGP with grow providing researchers with a tool to identify candidate genes for future experiments. MCRiceRepGP is available as a web application (http://mcgplannotator.com/MCRiceRepGP/)

**Significance statement:** Rice is a staple food crop plant for over half of the world’s population and sexual reproduction resulting in grain formation is a key process underpinning global food security. Despite considerable research efforts, much remains to be learned about the molecular mechanisms involved in rice sexual reproduction. We have developed MCRiceRepGP, a novel framework which allows prediction of sexual reproduction associated genes using multi-omics data, multicriteria decision analysis and machine learning. The genes identified and the methodology developed will become a significant resource for the plant research community.

## Introduction

Sexual reproduction is a core process in the life cycle of a vast majority of eukaryotic organisms. It is the main source of genetic diversity, which in turn allows for evolution and adaptation. From economic perspective, sexual reproduction results in formation of edible fruit and grains, underpinning crop yield and global food security. In plants, sexual reproduction is initiated by the vegetative to reproductive phase transition, requiring intricate signalling pathways integrating a multitude of internal and external cues. Upon commitment to flowering the process involves the development of reproductive organs, successful completion of male and female meiosis and fertilization, followed by embryonic development. Biological processes involved in sexual reproduction consist of evolutionarily conserved core components (for example, basic reproductive organ development including anthers and pistils and meiosis) (Schurko and Logsdon, 2008, Wallace *et al.*, 2011, Gómez *et al.*, 2015) and a species-specific regulatory level, for example the details of floral organ morphology and control of fine tuning of timing of vegetative to reproductive phase transition (Jarillo and Piñeiro, 2011, Moyroud and Glover, 2017). Knowledge of both, the level of conservation of core components and the species-specific characteristics of reproductive processes is crucial for understanding of plant fertility. Despite considerable research efforts the molecular basis of plant reproduction is not yet fully understood.

Rice is an important cereal crop, providing staple food for over a half of the world’s population. It is a monocotyledonous plant species with a relatively compact genome. The rice genome was one of the first plant genomes to be sequenced, providing a tremendous resource for plant research community. However, despite considerable research efforts, many of the genes involved in sexual reproduction remain uncharacterized (Kun *et al.*, 2013, Niu *et al.*, 2013, Fu *et al.*, 2014, Rhee and Mutwil, 2014, Hu *et al.*, 2015, Yao *et al.*, 2017). Several computational methods have been applied to improve understanding of gene functions. Studies of sequence homology between the most extensively studied and functionally annotated proteome of *Arabidopsis thaliana* and other species, including rice, allowed identification of genes with conserved functions (Gómez *et al.*, 2015). Construction of co-expression networks allowed identification of regulatory hubs involved in plant developmental processes, including anther development (You *et al.*, 2016, de Luis Balaguer *et al.*, 2017, Lin *et al.*, 2017). Analysis of expression profiles across tissues pinpointed genes with defined spatio-temporal expression patterns, which could be involved in organ, tissue or cell-specific processes (Edwards and Coruzzi, 1990). Studies of phenotypes of mutant lines provided annotation of genes with unknown functions (Miyao *et al.*, 2003, Miyao *et al.*, 2007). Genome-wide studies of diversity across hundreds of lines allowed identification of functionally important regions of increased or reduced diversity, helping pinpoint genes which display high sequence conservation within species (Alexandrov *et al.*, 2015, Tatarinova *et al.*, 2016).

Individually those approaches provide valuable insights into gene functions. The challenge is to combine all the resources into a unified framework to produce a list of reliable candidate genes involved in the biological process of interest (Troyanskaya *et al.*, 2003, Bradford *et al.*, 2010, Bargsten *et al.*, 2014). Our aim was to discover novel coding genes and lincRNAs involved in rice sexual reproduction. To achieve that we have developed a set of rules to prioritize the genes of interest and a novel method which combines information from gene expression studies, sequence homology, known functional annotation, mutational data and sequence diversity analysis. The method developed – MCRiceRepGP (Multi Criteria Rice Reproductive Gene Predictor) predicts gene’s potential for involvement is sexual reproduction using available multi-omics data, multi-criteria decision analysis, and machine learning. We applied the method to all rice genes and identified 2,275 protein coding and 748 lincRNA genes involved in rice reproductive processes. The manuscript also presents the first study of lincRNAs in plant gametes. A subset of the genes identified was linked to male and female-specific plant fertility. Several genes linked to reproductive stage heat stress tolerance were identified. For the purposes of the study, a full rice genome re-annotation using RNASeq datasets from 11 tissues and cell types has been performed. MCRiceRepGP is available as a web application (http://mcgplannotator.com/MCRiceRepGP/).

## Experimental procedures

### Datasets used

The rice genome assembly and annotation (MSU v7) and *A. thaliana* protein sequences (TAIR 10) were obtained from Phytozome v12.1 (Goodstein *et al.*, 2012). The RNASeq datasets were downloaded from Sequence Read Archive (Table S1). To maximize mapping specificity and minimize batch effects RNASeq from a minimum number of studies, covering maximum number of reproductive and vegetative tissues with read length equal or longer than 100 base were used. Phenotypic data for Tos17 rice mutant lines were downloaded from https://tos.nias.affrc.go.jp/ and the insertion coordinates were downloaded from http://orygenesdb.cirad.fr/. Gene ontology (GO) annotation of *A. thaliana* genes were downloaded from TAIR (ATH_GO_GOSLIM.txt, downloaded on: 20.07.2017) (Berardini *et al.*, 2015).

### Parameters used

Detailed commands for all the tools listed in the sections below can be found in Method S1.

### Genome reannotation

The RNASeq reads were mapped to the reference genome using Hisat2 v2.0.5 (Kim *et al.*, 2015) and the parameters were adjusted for stranded libraries. Transcripts were assembled separately for each library using StringTie v1.3.3b (Pertea *et al.*, 2015) and the parameters were adjusted for stranded libraries. The annotations were then merged with the existing rice annotation. lincRNAs were identified using procedure previously described (Golicz *et al.*, 2018b). In short, coding potential of genes was evaluated using Coding Potential Calculator 2 (Kang *et al.*, 2017) and homology to know protein coding genes. Transcripts were compared using DIAMOND v0.8.24.86 (Buchfink *et al.*, 2015) blastx against NCBI RefSeq (O’Leary *et al.*, 2016) protein database (downloaded on: 11.07.2017). A gene was considered coding if any of the transcripts were classified as coding by CPC2 or had a significant match in RefSeq database. The (long intergenic non-coding RNAs) lincRNAs were identified by comparing positions of coding and non-coding genes using bedtools (Quinlan and Hall, 2010). All non-coding genes which did not overlap any protein coding loci were classified as lincRNAs.

### Expression level evaluation

The reads mapping to gene loci were counted using featureCounts v1.5.1 (Liao *et al.*, 2014). The FPKM values were calculated as: (10^9^*fragments mapped to exons/assigned fragments*total length of exons). The log1p(FPKM) values were adjusted for batch effects using Combat v3.24.4 (Johnson *et al.*, 2007). The data used originated from three different studies, which was accounted for during batch effect adjustment.

### Homology analysis

For each coding gene representative (longest isoform) transcript was compared against the set of *A. thaliana* proteins (longest isoforms) using NCBI blastx v2.6.0 (Camacho *et al.*, 2009). GO annotations were transferred from *A. thaliana* genes to best matches (with lowest e-value) among the rice genes.

### Community analysis

Expression values were calculated by counting the number of reads mapping to each gene using FeatureCounts v1.5.1 (Liao *et al.*, 2014). The Spearman correlations were computed using corr function of psych package (Revelle, 2017). Top 5% of positive and negative correlations were used to built a co-expression network using Mutual Rank method (Obayashi *et al.*, 2009) (MR < 30). The Clique Percolation Method (Palla *et al.*, 2005) was used (https://sites.google.com/site/cliqueperccomp/) to identify putative functional modules within co-expression network. GO enrichment of nodes was calculated using topGO package v2.28.0 (Alexa *et al.*, 2006), using method ‘weight’ to adjust for multiple comparisons (p < 0.01).

### Diversity analysis

The filtered SNP set (18 M) was downloaded from SNP-Seek database (Alexandrov *et al.*, 2015). The number of SNPs falling within exons of each genes was counted and divided by total exon length of the gene as calculated by featureCounts. The gene was considered to be low diversity if the SNP density was below half of the median SNP density calculated using all genes.

### Process Involvement score parametrization

The Process Involvement (PI) score has seven components, which are weighted differently depending on their relative importance. Using knowledge of the field to supply probabilities for analysis of networks has been previously successfully applied (Troyanskaya *et al.*, 2003). The weights assume values between 0 and 1 and the values used were α=0.6, β=0.6, γ=0.4, δ=0.3, ε=0.2, ζ=0.1. The phenotypic data (P, α=0.6) and sequence homology with known sexual reproduction regulators (H, β=0.6) were considered to be the most important pieces of evidence and therefore were assigned the highest weight. Because one of the objectives of the study was to uncover key regulators of sexual reproduction, participation in functional co-expression modules was also considered important (CP, γ=0.4; CF, δ=0.3). Sequence diversity was also included, but given lower weighting. If genes had similar evidentiary support from phenotypic and/or homology data and network-connectivity, genes with lower diversity are hypothesized to be more likely regulators as transcription factors were shown to be the genes with lowest diversity in the rice genome (Tatarinova *et al.*, 2016). Finally, the expression value (EV, ζ=0.1) was given lowest weighting to prevent it from over-powering the entire score. Further details: Note S2.

### Classifiers

Three classifier were tested: (1) the Naïve Bayes classifier as implemented in function naiveBayes of package e1071 v1.6-8 (Meyer, 2017), (2) Classification Tree as implemented in function rpart of package rpart v4.1-11 (Therneau *et al.*, 2017), (3) Logistic Regression as implemented in method glm (R Core Team). Five-fold cross validation was used to measure the concordance between classifier prediction and test datasets. Further details: Method S2, Notes S3-S5.

### Test datasets

Ten genes known to have confirmed crucial roles in sexual reproduction were chosen as test dataset (Test Set 1) (Gómez *et al.*, 2015, Shi *et al.*, 2015a). Additionally, 781 genes implicated to be involved in sexual reproduction (https://funricegenes.github.io/) and highly expressed in reproductive tissues were used (Test Set 2).

### Fst score calculation

The 18M SNP dataset downloaded from SNP-Seek database was used. The *japonica* sub-population include temp and trop lines. The *indica* subpopulation included ind1, ind2 and ind3 lines. SNPs with minor allele frequency < 0.01 were remove from the dataset. Fst values were calculated using vcftools, with window size of 100kb and step of 10kb. Windows which fell within top 5% of highest Fst values (mean value) were retained, merged and compared with positions of SexRep genes.

### Data availability

Rice genome reannotation and files used as input for MCRiceRepGP can be found at: https://osf.io/78axs/.

Source code can be obtained from: https://github.com/agolicz/MCRiceRepGP and https://github.com/agolicz/MCRiceRepGP-shiny.

Web application can be found at: http://mcgplannotator.com/MCRiceRepGP/.

## Results

### Rice genome re-annotation using RNASeq data

The two available rice genome annotations (MSU-RAP and RAP-DB) were performed before RNASeq data was widely available and gene evidentiary support relied mostly on ESTs, which used to be derived from pools of samples, likely missing genes expressed at lower levels, transiently expressed or in low abundance cell types (Note S1). This is an especially important consideration while investigating sexual reproduction which depends on precise spatiotemporal gene expression regulating cell fate commitment and specification involving a small number of specialized cell types. Additionally, a mounting body of evidence accumulated since the last rice genome annotation points to important roles of long non-coding RNAs in sexual reproduction and those should also be included in the analyses (Golicz *et al.*, 2018a). Long intergenic non-coding RNA (lincRNA) annotation in rice has been performed previously (Zhang *et al.*, 2014, Wang *et al.*, 2015a), however the transcriptomes of egg, pollen sperm, and vegetative cells were not included.

Accordingly, we updated the MSU-RAP annotation using RNASeq data from multiple rice tissues and cell types (leaf, root, shoot, flower, seed, anther, pistils, sperm, cell, egg cell, vegetative cell). The final annotation comprised 56,118 loci, including 46,149 protein-coding and 9,969 lincRNA loci (Note S1, Table S2). The expression profile of newly discovered putative protein-coding loci (7,218 genes, 65.9% containing open reading frame (ORF) >100 amino acids and 42.4% containing complete ORF > 100 amino acids) was analysed, and 80.9% of genes showed highest expression levels in reproductive tissues, suggesting that a number of reproduction related genes may be missing from the available MSU-RAP annotation (Fig. S1). The updated annotation is well suited for the study of rice reproductive processes. It also highlights the significance of including expression data from specialized organs and low abundance cell types, especially those highly relevant to the study performed.

### The MCRiceRepGP method and its application for identification of reproduction associated genes in rice

Many publicly available rice genomic, transcriptomic and mutational datasets and databases exist (Ware *et al.*, 2002, Droc *et al.*, 2006, Miyao *et al.*, 2007, Alexandrov *et al.*, 2015, Wang *et al.*, 2015b). Using the updated genome annotation, these resources can be employed to help identify genes associated with biological processes of interest, in this case sexual reproduction. MCRiceRepGP uses information about seven features: tissue expression profile (tissue type and expression levels), connectivity within co-expression network, co-expression hub functional annotation, existing mutational data with phenotypic information, sequence homology and single nucleotide polymorphism (SNP) diversity to calculate gene score and predict whether the gene is involved in a biological process (Table 1, Fig 1).

**Table 1.** Features take 546 n into account when evaluating the PI score.

| Feature | General feature description | As applied for identification of sexual reproduction genes | Possible values | Parameter (feature weight) | Parameter values |
| --- | --- | --- | --- | --- | --- |
| Expression type (ET) | Is the highest expression recorded in a relevant tissue type? | Is the highest expression recorded in reproductive tissue? | 0 – no<br>1 – yes | $\alpha$ | 0.6 |
| Phenotype category (P) | Is the most common mutant phenotype consistent with the highest expression tissue type? | Is the most common phenotype associated with the transposable element insertion in the gene reproductive only? | 0 – no<br>1 – yes |  |  |
| Sequence homology (H) | Is the homologous *A. thaliana* gene annotated with relevant functions? | Is the homologous *A. thaliana* gene annotated with reproductive function? | 0 – no<br>1 – yes | $\beta$ | 0.6 |
| Community participation (CP) | Is the gene found within a community in co-expression network? | Is the gene found within a community in co-expression network? | 0 – no<br>1 – yes | $\gamma$ | 0.4 |
| Community function (CF) | Does the gene belong to a community annotated with relevant functions? | Does the gene belong to a community annotated with reproductive function? | 0 – no<br>1 – yes | $\delta$ | 0.3 |
| Sequence diversity (D) | Does the gene display low sequence diversity within the species? | Does the gene display low sequence diversity within the species? | 0 – no<br>1 – yes | $\varepsilon$ | 0.2 |
| Expression value (EV) | The FPKM value for the gene in the tissue with highest expression | The FPKM value for the gene in the tissue with highest expression | EV | $\zeta$ | 0.1 |

**Fig. 1.**
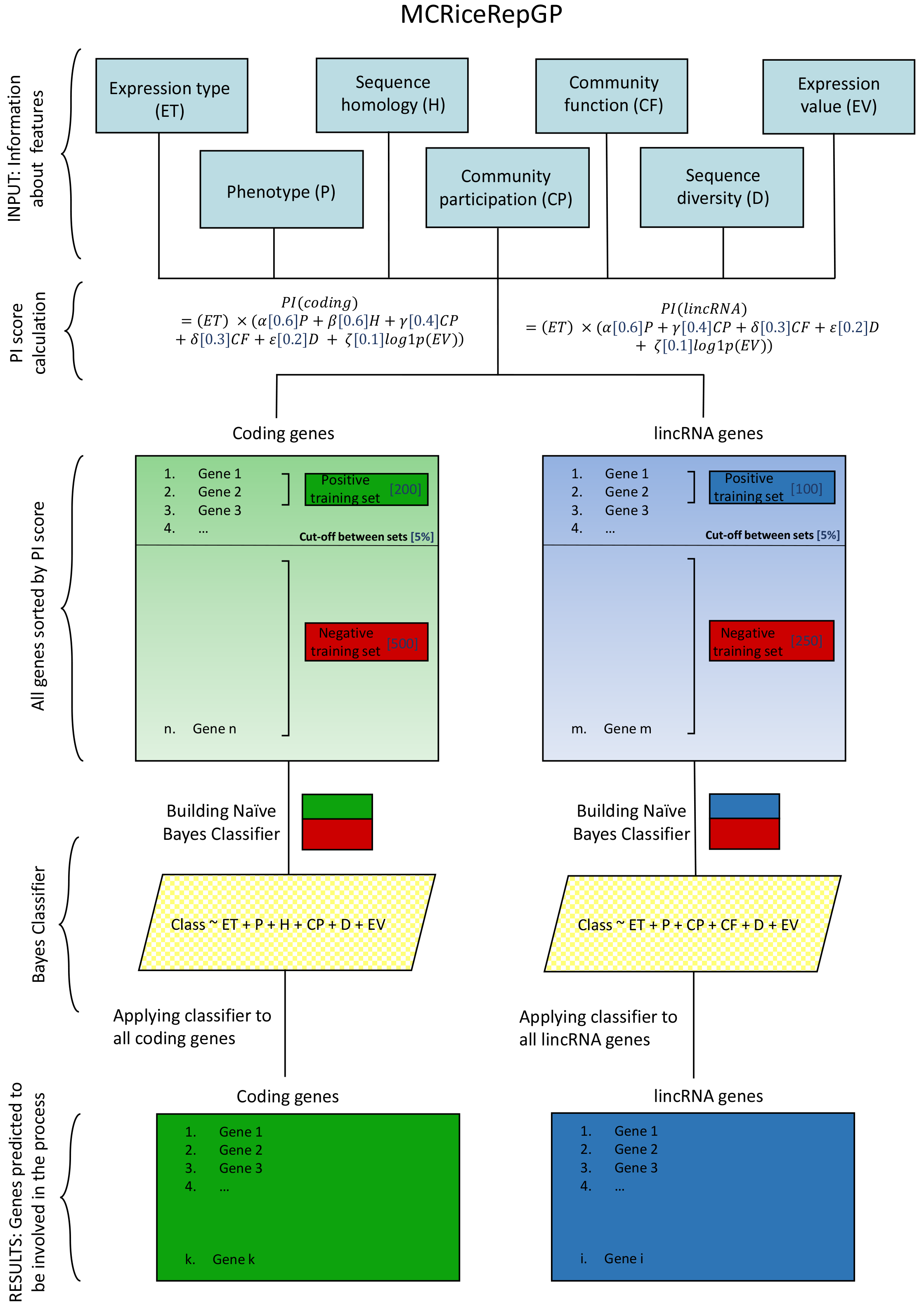
MCRiceRepGP method overview. Seven features (ET – expression type, P – phenotype category, H – sequence homology, CP – community participation, CF – community function, D – sequence diversity, EV – expression value) are used when evaluating a gene’s potential for involvement in a biological process. The features are scored and weighted and the Process Involvement (PI) score is calculated. The top and bottom scoring genes are used as positive and negative training set to build Naïve Bayes classifiers for coding an lincRNA genes. The classifiers are then used to identify a final set of genes involved in a given process. The values of parameters used to identify genes involved in sexual reproduction are presented in square brackets.

### Tissue expression profile analysis and gene co-expression network construction

#### Tissue expression profile analysis

Expression of all the rice genes across tissues was measured by quantifying number of RNASeq reads mapped to each gene locus and calculating FPKM (fragments per kilo base per million) value (Fig. S2). Because the dataset originated from several different studies, the expression values were adjusted in order to remove batch effects (Johnson *et al.*, 2007). Genes which are involved in a given process often show high or unique expression in related tissue (Wen *et al.*, 2016, Boyle *et al.*, 2017, Golicz *et al.*, 2018b). The samples were classified as either representing vegetative or reproductive tissue (Table S4). For each gene, the tissue and tissue type with highest expression levels observed was recorded. In total 72.6% (68.1% on non-batch-adjusted data) genes had the highest expression level in reproductive tissue/cell type. A high number of genes having peak expression in reproductive tissues is expected. Reproductive processes are complex, requiring developmental transitions, cell fate decisions and formation of multiple highly specialized cell types in male and female gametophytes, therefore are expected to engage a multitude of genes.

#### Co-expression network construction

The FPKM expression values were used to calculate all-vs-all Spearman correlations and the gene pairs within the top 5% (corresponding to minimum rho= 0.725 for positive correlations) or bottom 5% (corresponding to maximum rho= −0.619 for negative correlations) correlation values were used to build a co-expression network containing 50,212 nodes and 678,548 edges. Within the network, it is possible to identify sub-populations of tightly connected nodes – so called communities (Acharya *et al.*, 2012). These likely correspond to functional modules related to distinct biological roles. The whole network was analysed using Clique Percolation Method (Palla *et al.*, 2005), detecting 5,791 communities (putative functional modules). The modules were then functionally annotated using gene ontology (GO) enrichment analysis. Following the procedure used in the MSU-RAP annotation, the rice genes were annotated with GO terms corresponding to the most significant BLAST match in the *A. thaliana* proteome and GO enrichment for each module was calculated using all genes as background. The significantly enriched terms (p < 0.01) were assigned to modules as the functional annotation. In total, 4,044 modules were annotated with at least one GO term. The assigned terms were then manually inspected to identify key words/phrases associated with sexual reproduction (Table S5). Nodes which were annotated with at least one GO term containing a key word/phrase were annotated as associated with sexual reproduction (566 modules in total).

### Insertional mutant data

To date, the most comprehensive rice mutant panel with a published collection of phenotypes are the ~50,000 transposon Tos17 insertion lines (Miyao *et al.*, 2003, Miyao *et al.*, 2007). The link between disruption of gene sequence and the observed phenotype can be indicative of gene function. However, analysis of the dataset poses several challenges. Each line possesses more than one transposon insertion within the genome, with up to 10 Tos17 insertions per line (Miyao *et al.*, 2003). Not every insertion has a phenotypic manifestation, but in some cases, a single insertion can cause multiple aberrant phenotypes. In fact, almost half of the lines showed more than one phenotype (Miyao *et al.*, 2007). Because multiple Tos17 insertions within the genome of one line exist establishing a correlation between insertion and phenotype is not straight forward. However, if two or more lines have independent insertions in the same gene and exhibit the same/similar phenotype, disruption of the gene is likely linked to the phenotype. To facilitate detection of the most common phenotype associated with the insertion a more fuzzy match was performed – the 49 phenotypes were split into more general categories: reproductive timing, reproductive fertility, reproductive seed, reproductive organ, vegetative, lethal and dwarf (Table S6).

The insertion sites derived from all the lines were compared with exonic positions of genes. For each gene, all the lines which had an insertion within exons of the gene were extracted, and the most common phenotype and phenotype category (reproductive timing, reproductive fertility, reproductive seed, reproductive organ, vegetative, lethal and dwarf) were recorded. In total, 3,252 genes could be assigned at least one line with phenotype, and for 1,295 the most common phenotype was categorized as reproductive.

### Sequence homology analysis

Sexual reproduction is a process conserved in eukaryotes, with a number of genes involved in core processes, sharing sequence homology and conserved functions even among distantly related species (Schurko and Logsdon, 2008, Wallace *et al.*, 2011, Gómez *et al.*, 2015). For example, corresponding genes involved in anther and pollen development have been found (Gómez *et al.*, 2015). Therefore, the functionality of *A. thaliana* homologs can help in the prediction of roles of rice genes. The sequences of rice and *A. thaliana* genes were compared and GO annotation was transferred from *A. thaliana* genes to best rice gene matches. Additionally, the GO terms were compared with the list of key reproductive terms constructed during functional annotation of the co-expression network. Genes which were annotated with at least one GO term which contained a key word/term were annotated as associated with sexual reproduction.

### Sequence diversity analysis

Rice has the most extensive single nucleotide polymorphism database of any plants (Alexandrov *et al.*, 2015). The database lists ~20 million SNPs discovered using genomic data from ~3000 lines. Lower SNP density across genomic regions is associated with either purifying selection or selective sweeps (Wollstein and Stephan, 2015). An analysis of SNP diversity across the rice genome revealed that genes associated with regulation of transcription have lower than average sequence diversity (Tatarinova *et al.*, 2016) and transcription factor activity plays a key role in the control of biological processes. Furthermore, the known sexual reproduction master regulators (Table 2) were enriched in genes with low sequence diversity (Fisher exact, p < 0.05). The functional lncRNAs were also shown to have lower rates of evolution compared to non-functional ones (Wen *et al.*, 2016). Overall, 21.23% genes were identified as low diversity.

### Predicting gene’s potential for involvement in sexual reproduction

We devised a two-step approach in which we first apply Multi Criteria Decision Analysis (MCDA) based Process Involvement score (PI score) and then use the top scoring genes as the training dataset for Naïve Bayes classifier, which is in turn applied to the full set of genes. The combination of the classification provided by Naïve Bayes and the PI score ranking allows identification of most confident candidate genes involved in sexual reproduction.

#### Process Involvement (PI) gene score

The Process Involvement (PI) score is a single metric designed to measure gene’s potential for involvement in a biological process, in this case sexual reproduction. The score is inspired by Multi Criteria Decision Analysis (MCDA), a decision-making strategy used in a variety of settings from financial and urban planning to ecological risk assessment and medical diagnostics (DCLG, 2009, Adunlin *et al.*, 2015, Linkov *et al.*, 2015). MCDA involves combining multiple lines of evidence from different sources to aid complex problem solving. A general feature of MCDA is: 1. scoring of the options 2. weighting of the scores depending on their perceived importance. A similar approach can be used to evaluate the potential of gene’s involvement in biological process and prioritise genes with features of interest, given diverse evidentiary support including expression, sequence homology, and diversity data. (Fig. 1, Table 1).

Seven features are taken into consideration and combined to provide a single score. The score components were not weighted equally, ET, P and H contributing more to the score than CP, CF, D, and EV (Table 1, Experimental Procedures, Note S2). Overall, the genes which scored most favourably were: highly expressed in reproductive tissues, their disruption resulted in reproductive phenotype, had homologues in *A. thaliana* annotated with functions in reproduction, were highly connected in co-expression networks, had low sequence diversity among rice lines. The score for protein coding genes and lincRNAs differed slightly. For lincRNAs the homology term is ignored, as lincRNAs show little sequence conservation across species and very few have functional annotation. The PI score was calculated for all rice genes, resulting in a continuous distribution of scores (Fig. S3) and the genes were ordered by descending PI score. The highest ranking (top scoring) genes were considered to have a high potential for involvement in sexual reproduction.

#### Using top scoring PI genes as training dataset and choosing the optimal machine learning classifier

The high and low PI scoring coding and lincRNA genes can be then used as training data for a machine learning classification algorithm. The training dataset was composed of 200 coding and 100 lincRNA top scoring genes (as an example of genes involved in sexual reproduction–positive training dataset) and a random selection of 500 coding and 250 lincRNA genes from the bottom 95% of the ranking (as an example of genes not involved in sexual reproduction–negative training dataset). The GO enrichment analysis has shown the top 200 coding genes to be highly enriched in functions related to sexual reproduction (Table S7), while the selection of 500 genes from the bottom 95% showed no such enrichment (Table S8).

Three types of classifiers were tested (1) Naïve Bayes classifier, (2) Classification Tree, (3) Logistic Regression. A machine learning based classifier essentially performs the following task: ‘Given a set of genes A, find all the genes with similar properties in a larger set B.’ The classifiers were evaluated with respect to Matthews correlation coefficient (MCC), sensitivity and specificity (Fig 2a). Receiver operating characteristic (ROC) curves were also generated by plotting sensitivity against (1 – specificity) and the area under the curve (AUC) was compared (Fig 2a). To achieve a more balanced positive to negative set ratio, the negative training set was composed of randomly selected subset of a larger number of genes and the effect of the repeated selection on classifier performance was also tested (Fig 2a, Notes S3-S5). Overall, the Naïve Bayes classifier outperformed the other two other classifiers across all the metrics for both coding and lincRNA genes and was therefore chosen to perform the analysis (Fig 2b and Fig 2c). The superior performance of Naïve Bayes classifier for biological classification purposes using heterogenous data has been previously observed (Troyanskaya *et al.*, 2003, Bradford *et al.*, 2010, Sperschneider *et al.*, 2016). Additionally, Naïve Bayes classifier was shown to be not sensitive to the size of negative training set (Kurczab *et al.*, 2014, Kurczab and Bojarski, 2017) alleviating the potential effects of introducing artificial positive to negative training set ratio (Libbrecht and Noble, 2015).

**Fig. 2.**
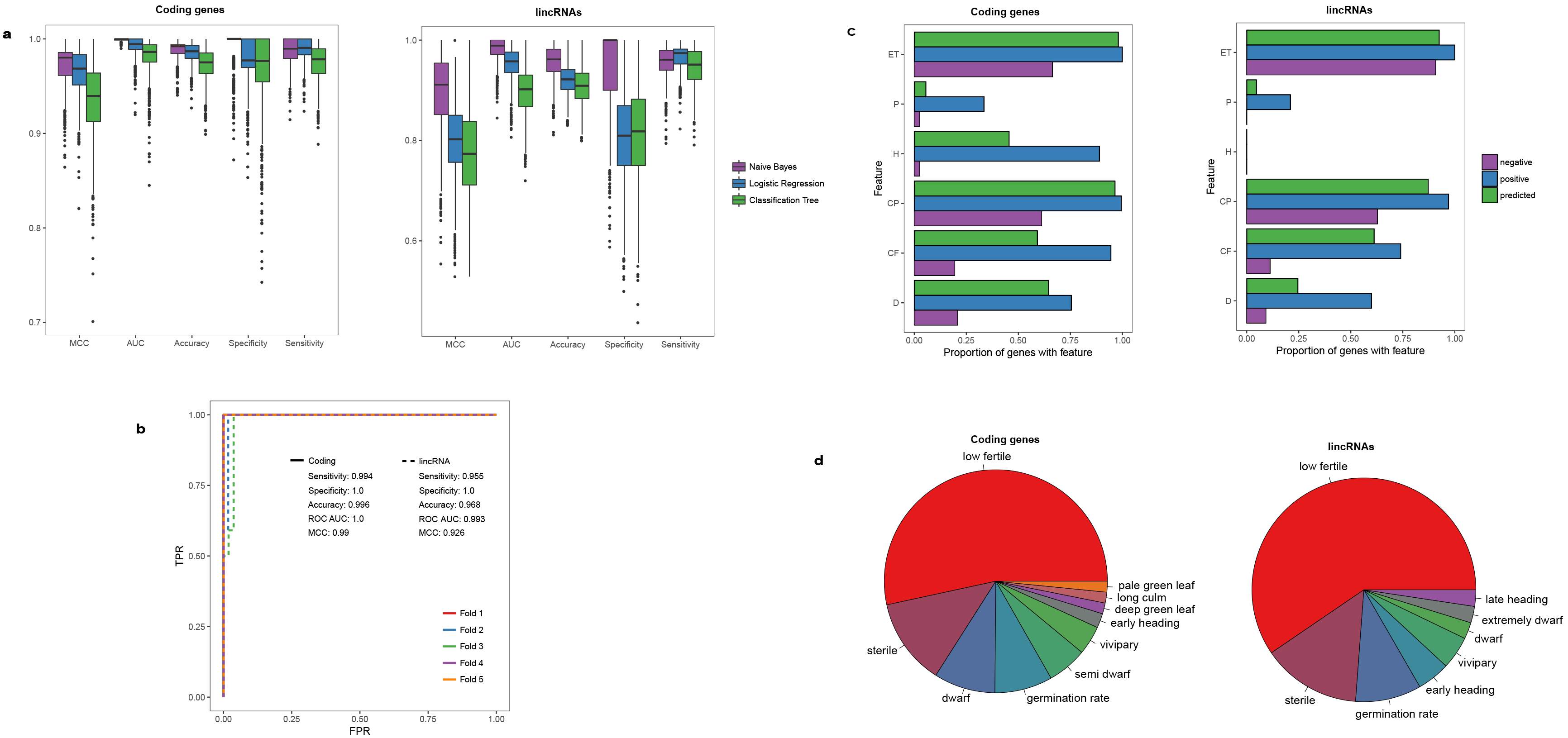
Comparison of the tested classifiers and the characteristics of the final Naïve Bayes classifier used for the analysis. (a) Three popular classifiers were tested: Naïve Bayes classifier, Classification Tree, and Logistic Regression. The performance measures used to assess the classifiers were: area under the receiver operating characteristic (ROC) curve (AUC) – interpreted as the ability of the classifier to distinguish between the two cases, MCC – Matthews correlation coefficient, sensitivity and specificity. The Naïve Bayes classifier was the top performing algorithm. The positive training sets were the top 200 and 100 coding and lincRNA genes as ranked by PI score. The negative training sets were the 500 and 250 genes randomly drawn from the bottom 95% of PI score gene ranking. In total, 200 negative training sets for coding and lincRNA genes were drawn and 5-fold cross validation for each negative set was performed (3×5×200 classifiers built for coding and lincRNAs genes). (b) The ROC curves along with other performance measures for the final Naïve Bayes classifiers for coding genes (200 top PI scoring coding genes as positive training set, random selection of 500 coding genes from the bottom 95% of PI score gene ranking as negative training set) and lincRNA genes (100 top PI scoring lincRNA genes as positive training set, random selection of 250 lincRNA genes from the bottom 95% of PI score gene ranking as negative training set). The performance of the classifiers was tested using 5-fold cross validation, the values provided are means. (c) Proportion of coding and lincRNA genes in positive training set, negative training set and final predicted SexRep genes which had value ‘1’ for the six binary features listed in Table 1 (ET – expression type, P – phenotype category, H – sequence homology, CP – community participation, CF – community function, D – sequence diversity). Vast majority of coding and lincRNA genes had peak expression in a reproductive tissue/cell type and belonged co-expression module(s). Many coding genes showed homology to known sexual reproduction regulators and had low sequence diversity. (d) Ten most common insertional mutant phenotypes for coding and lincRNA SexRep genes. The most common phenotype was low fertility and sterility.

#### Applying Naïve Bayes classifier

Naïve Bayes Classifier identified, 2,275 coding genes and 748 lincRNAs as involved in sexual reproduction (the genes identified by Naïve Bayes Classifier as involved in sexual reproduction were termed SexRep genes, Table S9). Again, GO analysis of SexRep genes showed strong enrichment of genes associated with sexual reproduction (Table S10). The number of genes involved in different reproduction related processes improved markedly when comparing the top 200 genes identified by PI score and the genes identified by Naïve Bayes classifier (for example, 54 and 347 genes respectively annotated as possibly involved in flower development; addition of 293 genes, addition of ~51 genes would be expected at random). The SexRep genes include 198 genes for which Tos17 mutant phenotypes were available (162 coding genes and 36 lincRNAs) (Fig. 2d). The four most common phenotypes were low fertile, sterile, germination rate and dwarf. This is consistent with observations that fertility and dwarf phenotypes are highly correlated (Miyao *et al.*, 2007).

#### Testing MCRiceRepGP predictions

The classifier has been trained to prioritize certain features including: high expression in reproductive tissues, homology to know *A. thaliana* proteins involved in reproduction and high connectivity in co-expression network. We have compared the results with a set of genes known to be crucial in rice sexual reproduction (Gómez *et al.*, 2015, Shi *et al.*, 2015a), which broadly fit into the criteria set while training the classifier (Test Set 1, Table 2). The genes represent a number of functional classes, including transcription factors, protein kinase, DNA de-methylase, Polycomb group protein and an lncRNA mi-RNA sponge (Nonomura *et al.*, 2003, Ono *et al.*, 2012, Yun *et al.*, 2013, Pan *et al.*, 2014, Wang *et al.*, 2017) and are involved in diverse processes including floral organ identity specification, floral patterning, sporogenesis, gamete fusion, endosperm and embryonic development. The method has classified all of those genes, including the lincRNA, as involved in reproduction. Additionally, we have tested the results against a database of 781 genes implicated to be involved in sexual reproduction (Test Set 2). Twenty eight percent of the Test Set 2 genes overlapped with SexRep genes and such an overlap is unlikely to occur by chance alone (permutation test, p < 0.01), confirming the suitability of the method for discovery of genes associated with sexual reproduction. Disregarding genes found in Test Sets 1 and 2 the method identified 2,060 coding and 747 lincRNA novel genes potentially involved in sexual reproduction.

**Table 2.** PI scores for known genes involved in sexual reproduction. Rep – predicted to be involved in sexual reproduction by MCRiceRepGP. MCRiceRepGP was not tested on LDMAR, a lincRNA known to be involved in rice sexual reproduction as it was not found in the annotation.

| Gene name | Gene ID | MSU-RAP Locus ID | Function | ET | P | H | CP | CF | D | Log1p(EV) | PI | MCRiceRepGP |
| --- | --- | --- | --- | --- | --- | --- | --- | --- | --- | --- | --- | --- |
| MADS3 | OSATST00001046 | LOC_Os01g10504 | Transcription factor | 1 | 0 | 1 | 0 | 0 | 1 | 5.16 | 1.316 | Rep |
| MADS58 | OSATST00036993 | LOC_Os05g11414 | Transcription factor | 1 | 0 | 1 | 1 | 1 | 0 | 4.85 | 1.785 | Rep |
| MADS15 | OSATST00045376 | LOC_Os07g01820 | Transcription factor | 1 | 0 | 1 | 1 | 0 | 0 | 3.68 | 1.368 | Rep |
| MADS1 | OSATST00025799 | LOC_Os03g11614 | Transcription factor | 1 | 0 | 1 | 0 | 0 | 1 | 5.21 | 1.321 | Rep |
| DL | OSATST00025795 | LOC_Os03g11600 | Transcription factor | 1 | 0 | 1 | 0 | 0 | 1 | 4.85 | 1.285 | Rep |
| MSP1 | OSATST00006904 | LOC_Os01g68870 | Kinase/Signalling | 1 | 1 | 1 | 1 | 1 | 1 | 2.41 | 2.341 | Rep |
| OsRos1a | OSATST00001213 | LOC_Os01g11900 | DNA demethylation | 1 | 0 | 1 | 1 | 0 | 1 | 3.71 | 1.571 | Rep |
| OsFIE2 | OSATST00049934 | LOC_Os08g04270 | Polycomb silencing | 1 | 0 | 1 | 1 | 1 | 0 | 3.54 | 1.654 | Rep |
| HAP2 | OSATST00037441 | LOC_Os05g18730 | Gamete fusion | 1 | 0 | 1 | 1 | 1 | 0 | 4.55 | 1.755 | Rep |
| Osa-eTM160 | NC_OSATST00025950 | N/A | miRNA sponge | 1 | 0 | N/A | 1 | 0 | 1 | 2.02 | 0.802 | Rep |

## Characterization of genes predicted to be involved in sexual reproduction

### Overall properties of genes predicted to be involved in sexual reproduction

The 3,023 SexRep genes (2,275 protein-coding genes and 748 lincRNAs) were analysed in more detail. Both coding and lincRNA SexRep genes showed an even distribution across chromosomes (Fig. 3a). The protein coding genes had higher overall expression levels when compared to lincRNAs, which is consistent with observations in rice and other plant species (Fig. 3b and Fig. 3c) (Zhang *et al.*, 2014, Wang *et al.*, 2015a). Analysis of tissue expression patterns of coding and lincRNA SexRep genes revealed that the highest proportion of genes had peak expression in egg and sperm cells respectively (Fig. 3b and Fig. 3c). Molecular function enrichment (Table S11) of the protein coding-genes showed them to be involved in protein binding, transcription factor activity, kinase activity and chromatin binding. Overall. 54.4% of the protein SexRep genes had no detectable similarity to *A thaliana* genes involved in sexual reproduction, but 59.2% were found in communities annotated with reproductive functions. Similarity, 61% of lincRNAs were found in communities annotated with reproductive functions (Fig 2c).

**Fig. 3.**
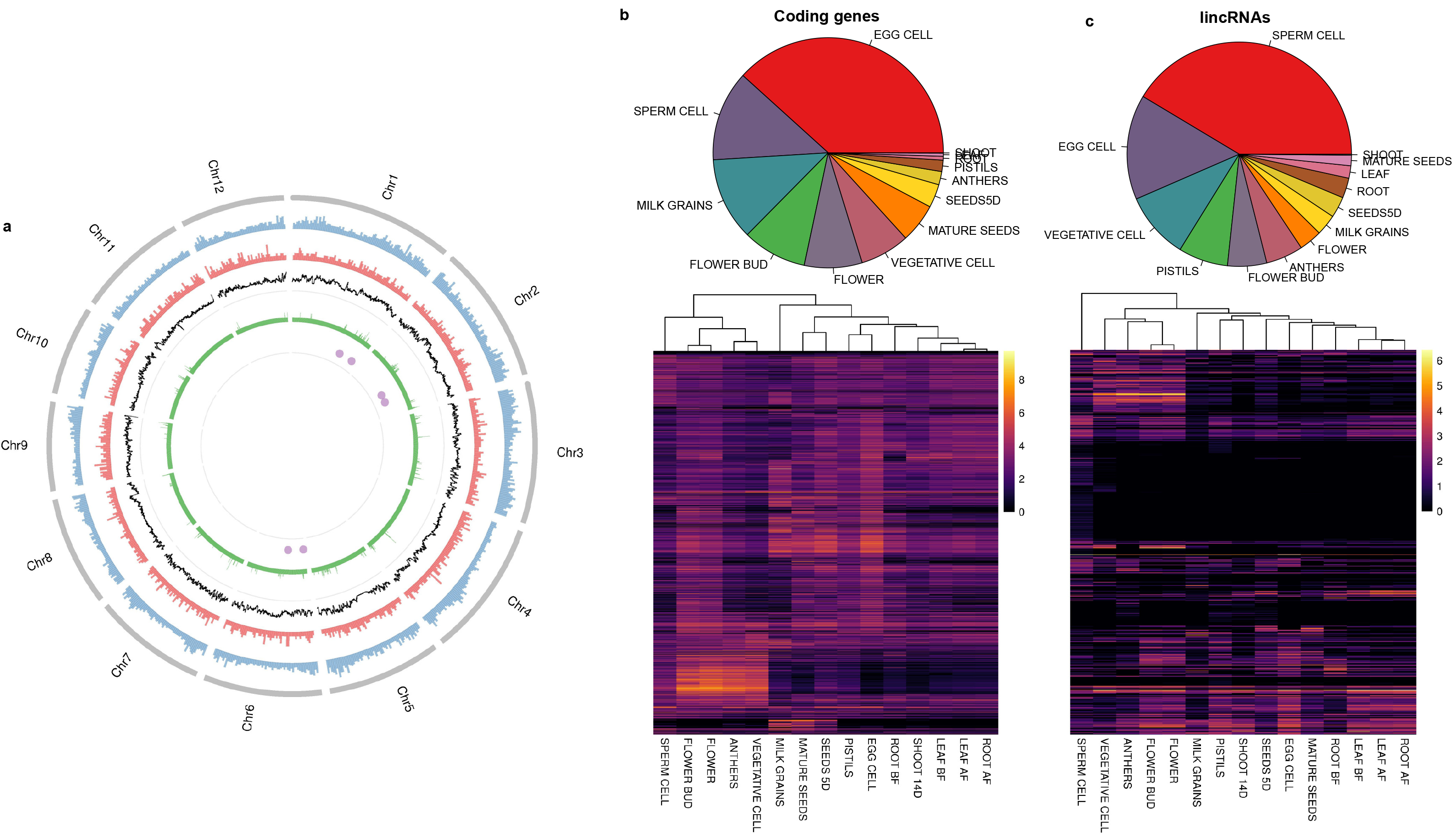
The landscape of SexRep genes. (a) Circular plot presenting SexRep gene distribution along the rice genome. From the outside ring: (1) coding SexRep genes, (2) lincRNA SexRep genes, (3) Fst index between *japonica* and *indica* sub-populations, calculated for 100 kb overlapping windows with a step of 10kb, (4) SexRep genes falling within regions of 5% highest Fst values (5) SexRep genes overlapping sterility associated loci identified in GWAS. (b,c) Heatmaps presenting expression of SexRep genes across tissues/cell types. Coding genes have higher overall expression values. Many of the lincRNA genes are expressed in sperm cells. Pie charts on top of heatmaps summarize the number of genes with peak expression in a given tissue/cell type.

**Table 3.**
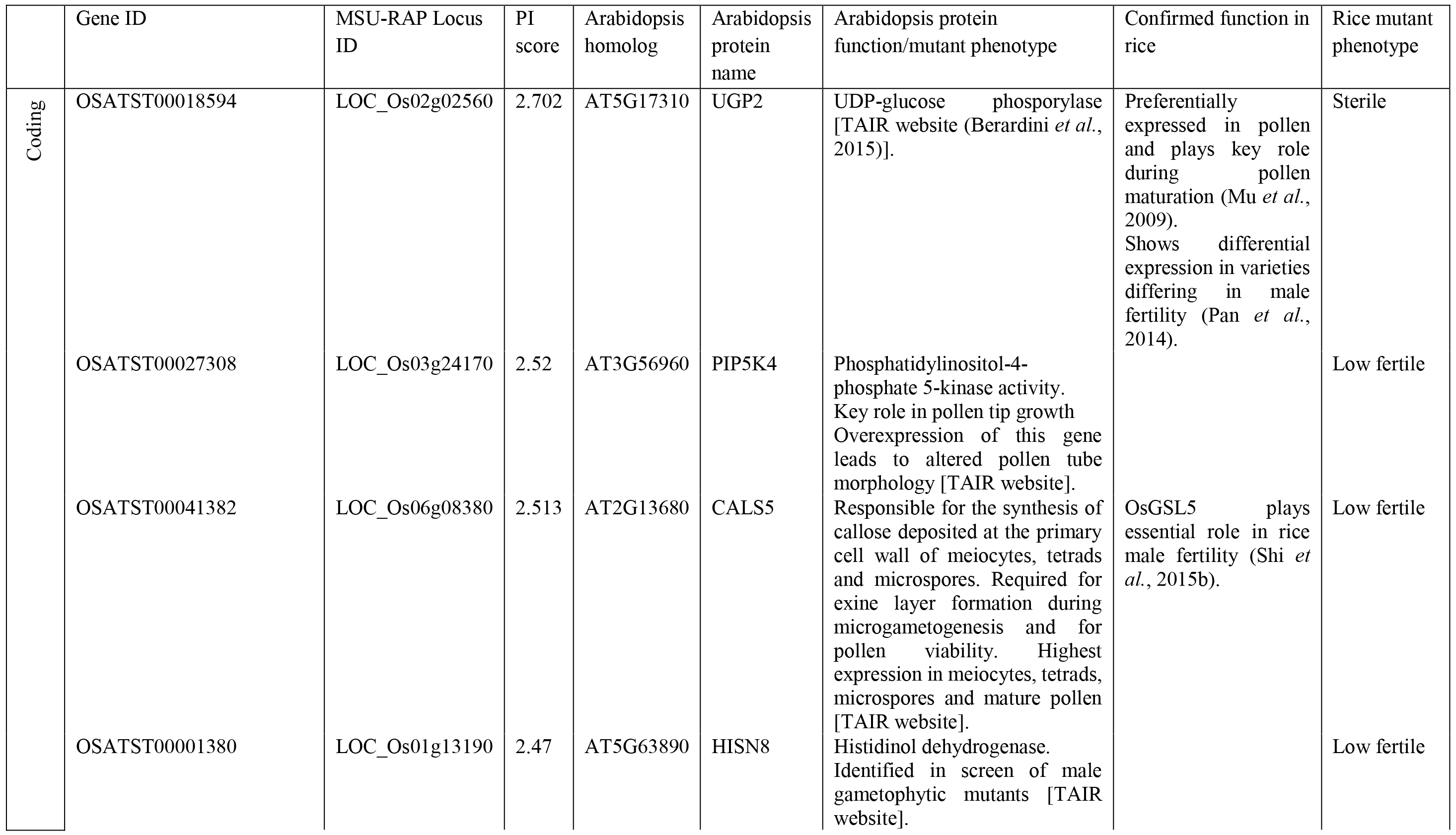
Top ten SexRep genes with highest PI scores.

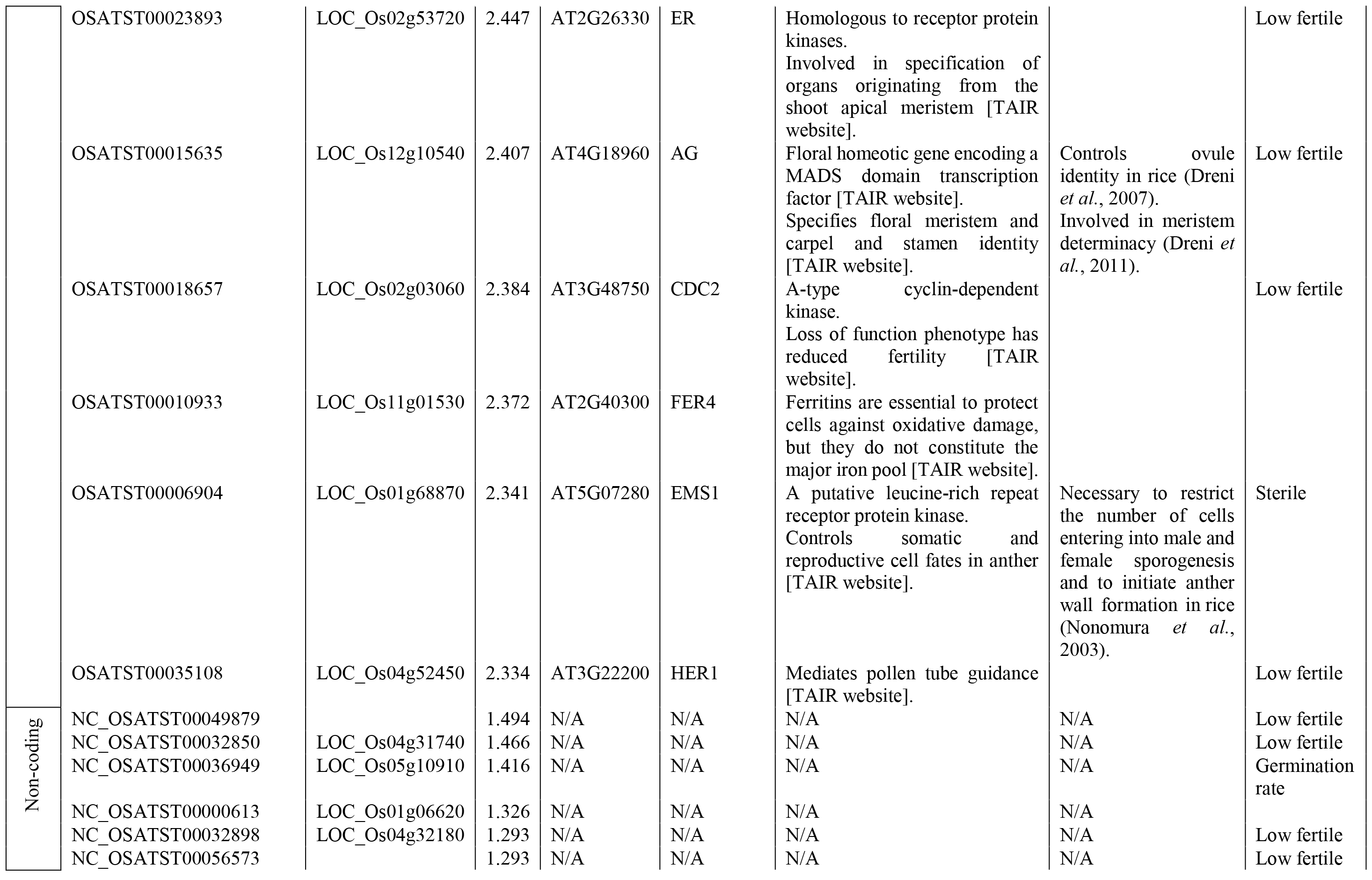

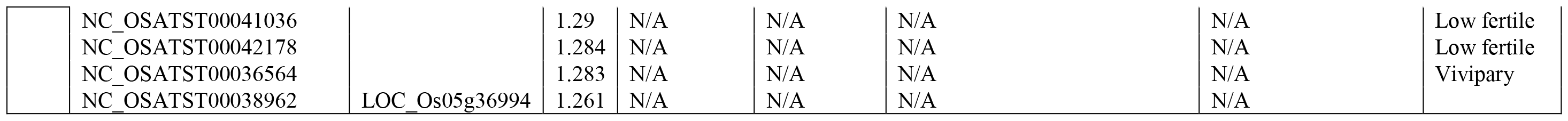

### Top candidate SexRep genes have diverse functional annotation

The SexRep genes can be ranked by PI score to identify most confident candidates. Top 10 SexRep genes (as ranked by PI score) were investigated in more detail (Table 3). Analysis of *A. thaliana* homologs suggests a diversity of molecular functions including protein kinases, transcription factor, UDP-glucose phosphorylase, histidinol dehydrogenase and ferritin. The genes appear to be involved in a range of processes from floral organ specification, cell cycle regulation, pollen maturation to pollen tube guidance. The most common phenotype found among the top ten genes was low fertility. To our knowledge four of the genes (LOC_Os01g68870, LOC_Os02g02560, LOC_Os06g08380 and LOC_Os12g10540) have already been characterized, confirming their involvement in sexual reproduction and influence on fertility (Yao *et al.*, 2017).

### SexRep genes have distinct tissue expression profiles

Genes which show unique or high activity in a given tissue are considered to be likely to contribute to the relevant biological processes (Wen *et al.*, 2016, Boyle *et al.*, 2017, Golicz *et al.*, 2018b). We investigated overall expression profiles of SexRep genes which show peak expression in a given tissue/cell type (Fig. 4a). Principal components analysis (PCA) shows clear clustering of both coding and lincRNA genes with peak expression in flower bud/flower, egg cells, pollen sperm cells and vegetative cells (Fig. 4a) suggesting that the genes may be involved in common biological processes. Protein coding SexRep genes have overall lower expression specificities (show broad expression across tissues/cell types), when compared to lincRNAs (lincRNAs are expressed in a limited number of tissues/cell types, Fig. 4b), which again is consistent with observations in other species (Golicz *et al.*, 2018a). For example, sperm cell SexRep protein-coding genes have one of the lowest median values of expression specificity index, while the lincRNA genes have the highest.

**Fig. 4.**
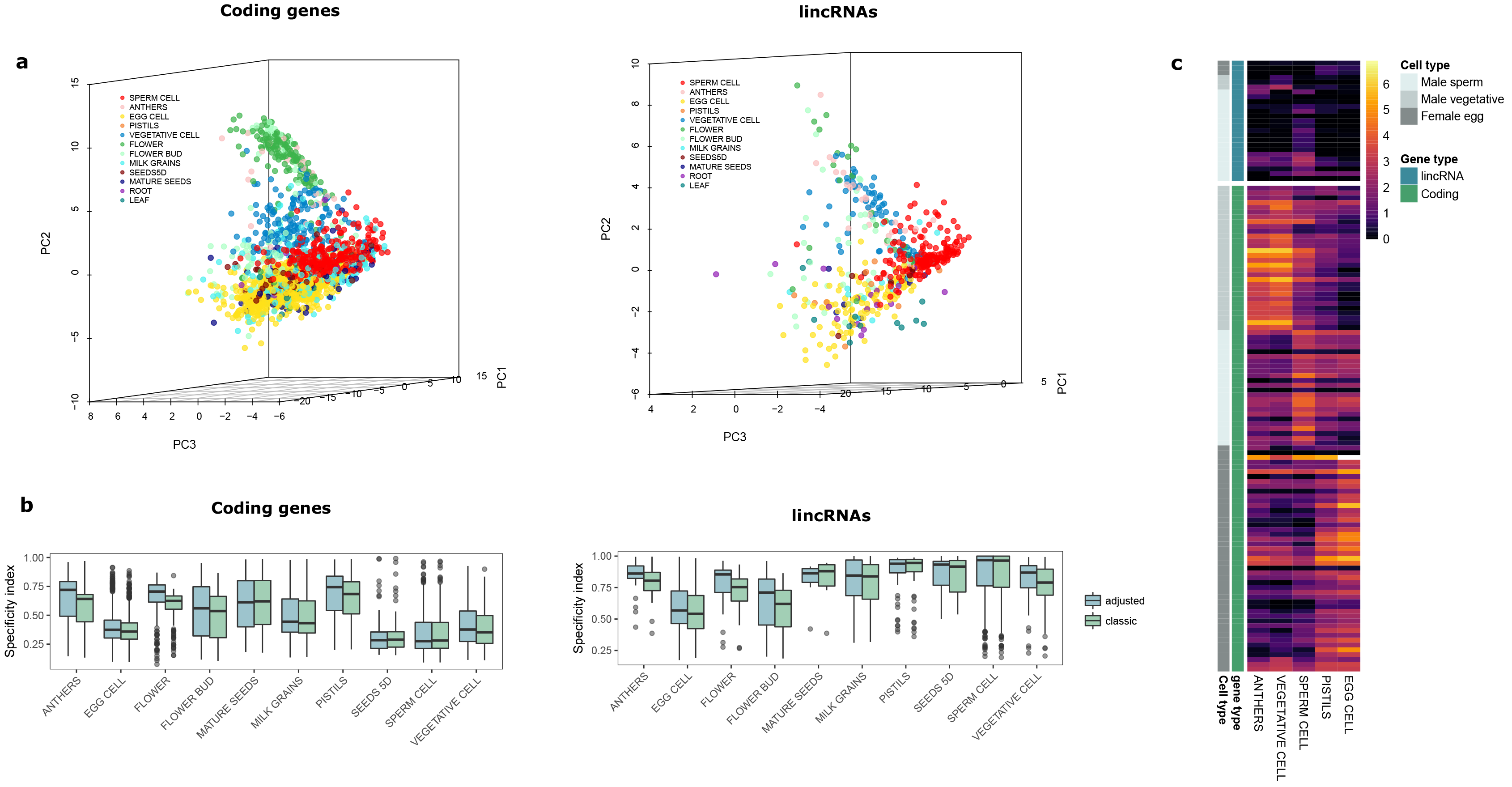
Overall expression patterns of SexRep genes with peak expression in a given tissue/cell type. (a) PCA analysis of coding and lincRNA gene expression values across tissues, each point corresponds to a gene and is coloured according to tissue/cell type in which the gene had peak expression value. Genes with common peak expression tissue/cell type cluster together – show similar overall expression patterns, which suggests involvement in common pathways/biological processes. (b) The box plots present tissue specificity index (*Tau*) of genes having highest expression point in a given tissue/cell type. Difference between coding and lincRNA genes can be observed. For example, the protein coding genes with peak expression in sperm cells have the lowest tissue expression specificities, while the lincRNA genes have the highest. The nested nature of sampling (for example, sperm cells are found within anthers, which in turn are found within flowers) could affect specificity calculations. Therefore, specificity indices were calculated twice, first using all samples (classic) and then adjusting for sample structure (adjusted). However, in both cases similar patterns were observed. (c) Heatmap presenting expression patterns of SexRep genes associated with fertility phenotype. The genes show sex-specific expression.

### Expression profile of SexRep genes suggests genes involved in male and female fertility

Sexual reproduction requires formation of reproductive structures including flower, anthers and pistils as well as successful male and female gametophyte development and fertilization. Defects which are sex specific will result in aberrant male or female fertility. We have investigated expression patterns of SexRep genes associated with fertility phenotype (Fig. 4c). Majority of the genes show sex-specific preferential expression. The genes associated with fertility phenotype show a clear split into three groups (1) genes with preferential expression in anthers and vegetative cells (2) genes with preferential expression in sperm cells and (3) genes with preferential expression in pistils and egg cells. Genes with preferential expression in male or female organs are potential contributors to sex-specific fertility.

### A subset of SexRep genes shows population differentiation between *japonica* and *indica* genotypes

In rice, there is an ancient and well-established divergence between two subspecies *japonica* and *indica* and the subpopulations are easily distinguishable based on their DNA sequence (Garris *et al.*, 2005). The subspecies also display phenotypic differences. For example, the *indica* lines being overall more heat tolerant than the *japonica* lines (Jagadish *et al.*, 2007, Zhao *et al.*, 2016), although heat tolerant lines exist in both sub-populations. Heat stress is known to reduce rice fertility with flowering (anthesis and fertilization) being the most susceptible stages of development (Jagadish *et al.*, 2007). The large polymorphism database available for rice (Alexandrov *et al.*, 2015) allows detailed genome-wide studies of differences between subspecies. The pairwise differentiation index (Fst) can be calculated between subpopulations, used to pinpoint regions of highest sequence diversity and find loci contributing to differences in phenotypes (Zhou *et al.*, 2015). In total, 288 SexRep protein coding genes fell within genomic regions corresponding to the top 5% of Fst values calculated between *japonica* and *indica* genotypes (Fig. 3a). GO enrichment analysis of those genes points to significant enrichment of genes associated with anther dehiscence (p = 0.0048, Table S12 and Table S13). Poor anther dehiscence is in turn known to be the leading cause of spikelet sterility induced by high temperatures due to poor efficiency of pollen delivery to stigma (Jagadish *et al.*, 2010, Zhao *et al.*, 2016). High differentiation of anther dehiscence related genes is consistent with observations of differential heat tolerance of *indica* and *japonica* sub-species.

### Several SexRep genes overlap loci associated with sterility in rice

The method used for detection of SexRep genes can also be used to enhance findings of genome wide association studies (GWAS). GWAS have been successfully used to uncover genomic regions containing loci associated with agronomic traits (Huang *et al.*, 2010, Yano *et al.*, 2016). Although high density SNP maps give good resolution to GWAS studies, usually several candidate genes within the region of interest are identified (Dingkuhn *et al.*, 2017). Usage of additional lines of evidence such as the ones used for identification of SexRep genes can help point to more confident candidates within the sections of the genome identified by GWAS. We have compared the genomic locations of recently identified SNPs linked to heat stress associated sterility in rice (Dingkuhn *et al.*, 2017) with coordinates of SexRep genes and identified six genes potentially related to sterility (Table S14). The number of SexRep genes found in vicinity of sterility associated SNPs (closer to the SNP than any other gene) was higher than it would be expected by chance (Chi Square, p < 0.01).

## Discussion

Despite considerable research efforts genes controlling sexual reproduction in plants remain enigmatic. Computational biology approaches can provide new insights by combining and analysing large-scale data from a number of sources, including genomic, transcriptomic and mutational datasets. The main challenge is the effective integration of all the information available. In this study, Process Involvement (PI) score and Naïve Bayes Classifier were applied to identify genes involved in sexual reproduction. MCRiceRepGP depends on seven features which describe the gene in terms of expression profile, biological network connectivity, homology with known sexual reproduction regulators and overall sequence diversity. MCRiceRepGP was applied to protein coding genes as well as non-coding RNA loci and identified three thousand protein coding genes and lincRNA loci involved in sexual reproduction. Analysis of all protein coding genes predicted to be involved in sexual reproduction (SexRep genes) highlighted genes involved in protein binding, transcription factor and kinase activity. The most common mutant phenotype associated with both coding and lincRNA SexRep genes was low fertility. The top SexRep protein coding genes had diverse functional annotations and are implicated in processes from floral organ specification, pollen development to pollen tube guidance. The genes identified are valuable resource providing potential targets for further experiments, including many long non-coding RNAs. Previous studies have shown long non-coding RNAs to play active roles in reproductive processes and the candidates identified in this study can open new avenues for rice research.

In this analysis MCRiceRepGP was parametrized to favour genes highly or specifically expressed in reproductive organs and with sequence homology to *A. thaliana* genes. However, alternative parameters can be chosen depending on the experimental goals. Other mutant lines can also be utilized. Recently a comprehensive library of neutron mutants became available, although no phenotypes have yet been recorded (Li *et al.*, 2017). Additionally, looking at individual components of the PI score can also point to genes of interest. For instance, looking only at genes which do not have homologs in *A. thaliana*, could help uncover rice specific regulators.

The method can be used in conjunction with other genome wide analyses. A number of genome-wide screens which help identify genomic regions associated with traits exist. These include genome-wide association studies (GWAS) to identify loci linked to traits of interest or calculation of fixation index (Fst) between sub-populations and identification of genomic regions with high and low differentiation. However, regions identified usually contain multiple genes, and it is not clear which one affects the trait. For example, in a recent GWAS study genes within ± 100kb of associated polymorphism were considered (Dingkuhn *et al.*, 2017) and the Fst values are also calculated for ~100kb windows (Zhou *et al.*, 2015). Often sequence homology only is used, but combining multiple lines of evidence can give more confident candidate gene predictions. Comparison of genome coordinates of SexRep genes against regions of high differentiation between *japonica* and *indica* genotypes revealed overrepresentation of genes associated with pollen release from anther, while comparison with GWAS data identified six genes potentially related to sterility.

A web application which implements MCRiceRepGP has been made available (Fig. 5). The application allows building of new classifiers by varying of PI score parameters, key words and classifier features. The results can be browsed online and are available for download.

**Fig. 5.**
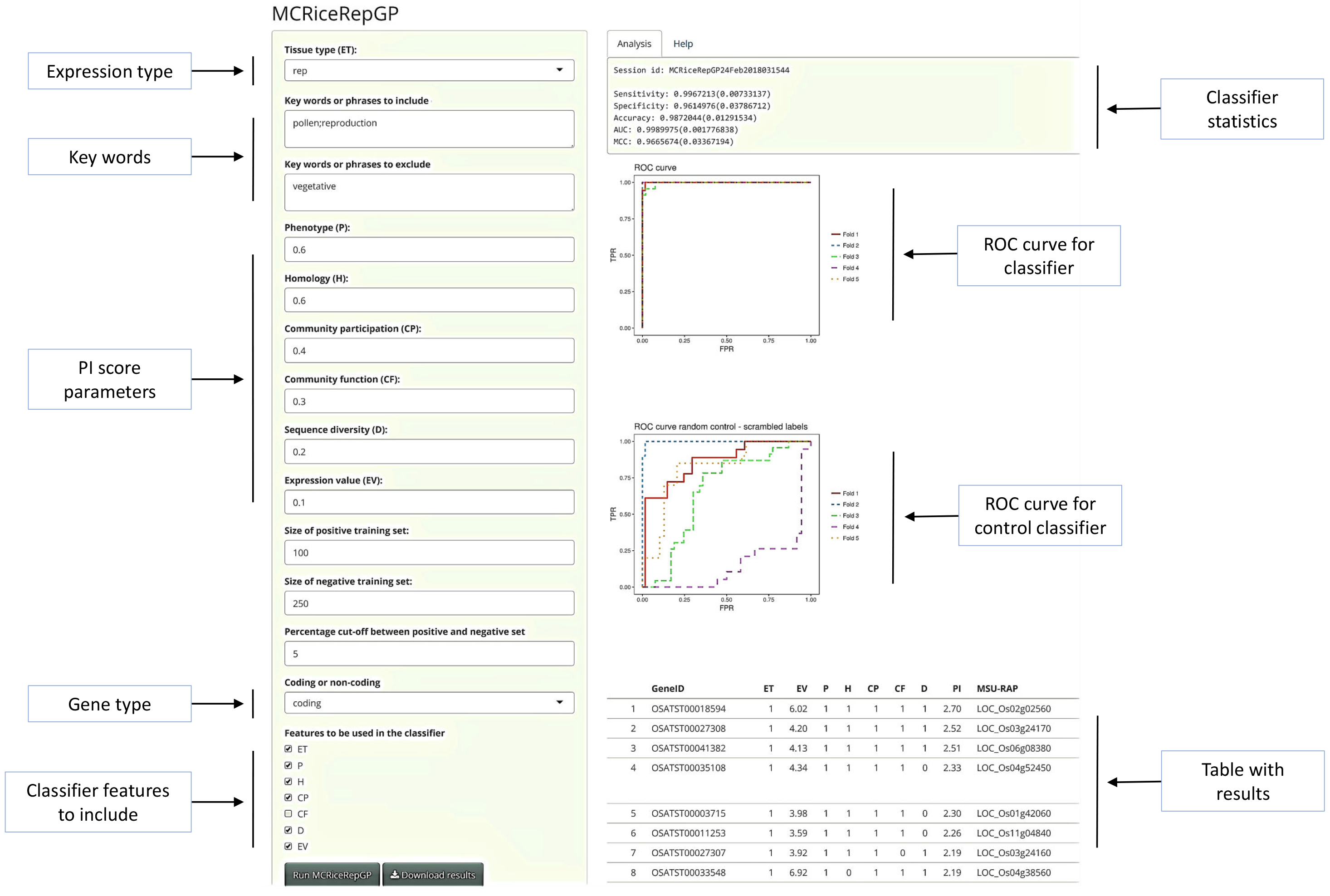
Screen shot of MCRiceRepGP web app results. The panel on the left side allows the user to control gene type, key words, PI score parameters and features to be included in the classifier. Results are displayed on the right-side panel. Results include classifier statistics, classifier ROC curve, control classifier ROC curve and the table with the final results.

## Conclusion

We have developed MCRiceRepGP – a method which combines evidence from heterogenous data sources for identification of novel genes involved in rice sexual reproduction. An easy to use web application has been made available and allows building of different classifiers. Additionally, for this study an updated rice genome annotation has been generated using deep sequencing data from reproductive tissues and cell types. The methodology developed, the putative reproduction associated genes and especially lincRNAs identified using MCRiceRepGP as well as the new rice genome annotation provide a valuable resource for further studies of rice sexual reproduction. Identification of previously unannotated genes from sexual reproduction specific tissues highlights the importance of including expression data from specialized organs and low abundance cell types in the genome annotation efforts. The novel sexual reproduction associated genes and lincRNAs described in the study provide targets for future research efforts. The method described may become an inspiration and an example of how different types of data can be integrated to predict most confident candidate genes and future research targets.

## Acknowledgements

This research was supported by Melbourne Bioinformatics at the University of Melbourne, project UOM0033. The research was supported by ARC Discovery grant DP0988972 and the University of Melbourne McKenzie Postdoctoral Fellowship.

## Authors’ contributions

AAG designed and performed the experiments, wrote the manuscript. MBS conceived research, designed the experiments, wrote the manuscript. PLB conceived research.

## Supporting Information

**Fig. S1** Distribution of tissues/cell types with peak expression levels of putative protein coding genes not found in MSU-RAP annotation

**Fig. S2** Heatmap representing expression of all coding and non-coding genes

**Fig. S3** Distribution of PI scores for coding and non-coding genes

**Fig. S4** Comparison of classifier performance for three different biological processes

**Fig. S5** Comparison of classifier performance for three different biological processes with scrambled labels

**Fig. S6** Number of shared SexRep genes identified by Naïve Bayes Classifier while varying alpha and zeta parameters of PI score

**Fig. S7** Overlap between results of five MCRiceRepGP runs using different negative training sets for coding and lincRNA genes

**Table S1** Datasets used in the analysis

**Table S2** Summary of annotation statistics

**Table S3** Comparison between current annotation and existing lincRNA annotations

**Table S4** Classification of samples as reproductive or vegetative

**Table S5** Dictionary of GO phrases associated with sexual reproduction

**Table S6** Classification of phenotypes as vegetative or reproductive

**Table S7** Biological process GO enrichment of 200 top genes as ranked by PI score

**Table S8** Biological process GO enrichment of 500 randomly chosen genes from bottom 95% as ranked by PI score

**Table S9** All SexRep genes identified

**Table S10** Biological processes GO enrichment of SexRep genes

**Table S11** Molecular function GO enrichment of SexRep genes

**Table S12** Biological processes GO enrichment of SexRep genes falling within top 5% of most differentiated genomic regions between indica and japonica lines

**Table S13** Anther dehiscence associated genes found in Supplementary table 12

**Table S14** SexRep gene overlapping sterility associated SNPs Method S1 Commands used for external software packages Method S2 Details of classifier implementation

**Note S1** Rice genome re-annotation

**Note S2** PI score parametrization for sexual reproduction

**Note S3** Comparison of classifiers used for identification of genes involved in three distinct biological processes

**Note S4** Naïve Bayes Classifier Performance with varying Process Involvement score parameters

**Note S5** Concordance between genes identified by classifiers built using different negative training sets

